# Genetic interactions among ADAMTS metalloproteases and basement membrane molecules in cell migration in *Caenorhabditis elegans*

**DOI:** 10.1101/2020.09.30.320093

**Authors:** Ayaka Imanishi, Yuma Aoki, Masaki Kakehi, Shunsuke Mori, Tomomi Takano, Yukihiko Kubota, Hon-Song Kim, Yukimasa Shibata, Kiyoji Nishiwaki

**Affiliations:** Department of Bioscience, Kwansei Gakuin University, Sanda, Japan

## Abstract

During development of the *Caenorhabditis elegans* gonad, the gonadal leader cells, called distal tip cells (DTCs), migrate in a U-shaped pattern to form the U-shaped gonad arms. The ADAMTS (<u>a</u> <u>d</u>isintegrin <u>a</u>nd <u>m</u>etalloprotease with <u>t</u>hrombo<u>s</u>pondin motifs) family metalloproteases MIG-17 and GON-1 are required for correct DTC migration. Mutations in *mig-17* result in misshapen gonads due to the misdirected DTC migration, and mutations in *gon-1* result in shortened and swollen gonads due to the premature termination of DTC migration. Although the phenotypes shown by *mig-17* and *gon-1* mutants are very different from one another, mutations that result in amino acid substitutions in the same basement membrane protein genes, *emb-9/collagen IV a1, let-2*/*collagen IV a2* and *fbl-1/fibulin-1*, were identified as genetic suppressors of *mig-17* and *gon-1* mutants. To understand the roles shared by these two proteases, we examined the effects of the *mig-17* suppressors on *gon-1* and the effects of the *gon-1* suppressors and enhancers on *mig-17* gonadal defects. Some of the *emb-9, let-2* and *fbl-1* mutations suppressed both *mig-17* and *gon-1*, whereas others acted only on *mig-17* or *gon-1*. These results suggest that *mig-17* and *gon-1* have their specific functions as well as functions commonly shared between them for gonad formation. The levels of collagen IV accumulation in the DTC basement membrane were significantly higher in the *gon-1* mutants as compared with wild type and were reduced to the wild-type levels when combined with suppressor mutations, but not with enhancer mutations, suggesting that the ability to reduce collagen IV levels is important for *gon-1* suppression.

## Introduction

Members of the ADAMTS family of secreted zinc metalloproteases have important roles in animal development. Most of these proteases degrade extracellular matrix components such as proteoglycans or collagens [1]. Nineteen ADAMTS genes have been identified in the human genome, and mutations in many of them result in hereditary diseases that are related to disorders of the extracellular matrix [2, 3]. Multiple ADAMTS proteases often function in a common developmental process. For example, the functions of ADAMTS-5, -9 and -20 are required for interdigital web regression [4]. ADAMTS-9 and -20 are needed for closure of the palate and for craniofacial morphogenesis and neural tube closure through their function in ciliogenesis [4, 5]. ADAMTS-5 and -15 act in myoblast fusion [6]. These enzymes appear to function in a partially overlapping manner. However, the roles shared by these ADAMTS proteases in development still remain elusive.

Among five ADAMTS genes in *C. elegans, gon-1* and *mig-17* play essential roles in the development of the somatic gonad [7, 8]. Both GON-1 and MIG-17 localize to the gonadal basement membrane (BM) [9, 10]. In the *mig-17* mutants, DTCs meander and stray, resulting in an abnormal gonadal shape. In contrast, in the *gon-1* mutants, DTCs rarely migrate and the gonads remain small. We have isolated and analyzed the genetic suppressors for DTC migration defects in *mig-17* mutants. Dominant gain-of-function mutations in the *fbl-1/fibulin-1* and *let-2/collagen IV α2* chain, which encode BM molecules, have been frequently isolated as suppressors [11, 12]. The *fbl-1(gf)* mutations are amino acid substitutions in the second epidermal growth factor–like motif of FBL-1C, one of the two splicing isoforms, and FBL-1C protein is secreted by the intestine to be released into the gonadal BM in a MIG-17 activity-dependent manner [11]. Suppression of *mig-17* by *fbl-1(gf)* mutations is dependent on the BM molecule NID-1/nidogen [12]. *let-2(gf)* mutations are associated with amino acid changes within the triple helix domain or the C-terminal non-collagenous domain (NC)1 of collagen IV. LET-2 protein is secreted from body wall muscle cells and DTCs and localized to the gonadal BM in a MIG-17 activity-independent manner. Unlike the case of *fbl-1(gf)*, suppression of mig-17 by *let-2(gf)* mutations does not require NID-1 [12].

Genetic suppressor analysis of *gon-1* mutants revealed that loss-of-function (deletion) mutations in *fbl-1* can suppress the shortened-gonad phenotype of *gon-1* mutants. Because *fbl-1* deletion mutants also exhibit the shortened-gonad phenotype, GON-1 and FBL-1 act antagonistically to regulate gonad formation [13]. This genetic interaction is mediated by the control of collagen IV accumulation in the gonadal BM: GON-1 acts to reduce collagen IV levels, whereas FBL-1 acts to maintain collagen IV levels [14].

Although *mig-17* and *gon-1* mutants are phenotypically very different, they are both suppressed or enhanced by mutations in *let-2* and *fbl-1*. In this study, we isolated novel suppressor mutations in *emb-9* for *mig-17* and in *let-2* for *gon-1* gonadal defects. Together with the previously isolated suppressors and enhancers, we investigated the consequences when suppressors for *mig-17* were combined with *gon-1* mutations, and when *gon-1* suppressors and enhancers were combined with *mig-17* mutations. We found that some of the *emb-9, let-2* and *fbl-1* mutations suppressed both *mig-17* and *gon-1*, whereas others suppressed only *mig-17* or *gon-1*. These results suggest that *mig-17* and *gon-1* have their specific functions as well as the functions commonly shared between them for gonad formation.

## Materials and Methods

### Strains and genetic analysis

Culture, handling and ethyl methanesulfonate (EMS) mutagenesis of *C. elegans* were conducted as described [15]. The following mutations and transgenes were used in this work: *mig-17(k174), gon-1(e1254, g518), fbl-1(k201, tk45), let-2(g25, b246, k193, k196), emb-9(tk75, g34, g23cg46)* and *tkTi1[emb-9::mCherry]* [7, 8, 11, 12, 14, 16-18]. The suppressor mutations were genetically mapped with single-nucleotide polymorphism mapping using *mig-17(k174)* and *gon-1(e1254)* mutant strains, which are in the CB4856 background [19]. Among the 11 *mig-17(k174)* suppressors, *k204* and *k207* were mapped to the center of linkage group (LG) III. Next-generation sequence analysis identified missense mutations in *emb-9* in these suppressors. Of the two *gon-1(e1254)* suppressors, one was mapped to the right end of LG X and was identified by next-generation sequence analysis as corresponding to a missense mutation in *let-2*. All experiments were conducted at 20°C. The temperature-sensitive mutants *emb-9(g34)* and *let-2(g25, b246)*, which arrest during embryogenesis or early larval stages at 25°C, do proliferate at 20°C. Because *gon-1(e1254, g518)* and *fbl-1(tk45)* mutants were sterile, we used the genetic balancer *nT1[qIs51] (IV;V)* to maintain these mutants and to generate double mutants containing these mutations.

### Microscopy

Gonad migration phenotypes were scored using a Nomarski microscope (Axioplan 2; Zeiss). Analysis of gonadal phenotypes was performed at the young-adult stage as described [20]. Although the gonadal phenotypes of some strains used are published, we reevaluated them in this study. The levels of EMB-9-mCherry were quantified as follows. For each sample, confocal images of a sagittal section of the DTCs were obtained with a Zeiss Imager M2 microscope equipped with a spinning-disk confocal scan head (CSU-X1; Yokogawa) and an ImageEM CCD camera (ImageEM; Hamamatsu Photonics). Using ImageJ software, we measured the fluorescence intensities along three drawn lines, each of which crossed the DTC BM; the average background intensities inside the gonad were subtracted from the peak values of the line scan, and the resulting corrected values were averaged.

### Transgenic analysis of suppressors

Germline transformation was carried out as described [21]. Plasmids containing *emb-9(k204), emb-9(k207)* and *let-2(tk101)* were constructed by introducing these mutations individually into their respective wild-type constructs [12, 14]. These plasmids were injected into the *unc-119(e2498)* gonad at 1–2 ng/μl with 10 ng/μl *unc-119+* plasmid [22], 70 ng/μl *sur-5::gfp* plasmid [23] and 70 ng/μl pBSIIKS(–), and the resulting extrachromosomal arrays were transferred to either *mig-17(k174); unc-119(e2498)* or *gon-1(e1254); unc-119(e2498)* animals by mating.

### Homology modeling of NC1 domains

Homology modeling was conducted based on the crystal structures of bovine collagen IV NC1 domains [24] using the SWISS-MODEL server. Ribbon diagrams were created and edited with the Swiss-Pdb Viewer software.

## Results

### Isolation of novel suppressors of *mig-17* and *gon-1* mutants

The *C. elegans* hermaphrodite gonad arms extend to the anterior-right and posterior-left areas of the body cavity. The U shape of the gonad arms are generated by migration of the gonadal leader cells, the DTCs, over the body wall during larval development (S1 Fig). *gon-1* mutants exhibit shortened gonads due to the premature termination of DTC migration and are sterile. In contrast, *mig-17* mutants show misshapen gonad arms due to the misdirected migration of DTCs, but they are still fertile. Although both of these genes encode ADAMTS family metalloproteases, the phenotypes shown by these mutants are very different.

The null allele *gon-1(q518)* exhibits a fully penetrant short gonad phenotype that cannot be distinguished from that of the *mig-17(k174)*; *gon-1(q518)* double null mutants [8]. The phenotypic penetrance of *gon-1(e1254)*, a partial loss-of-function allele that results in a milder gonad phenotype as compared with *gon-1(q518)*, was enhanced in combination with the *mig-17(k174)* null allele (Fig 1A; S2 Fig), indicating that *gon-1* and *mig-17* function in partially overlapping pathways.

**Fig 1.**
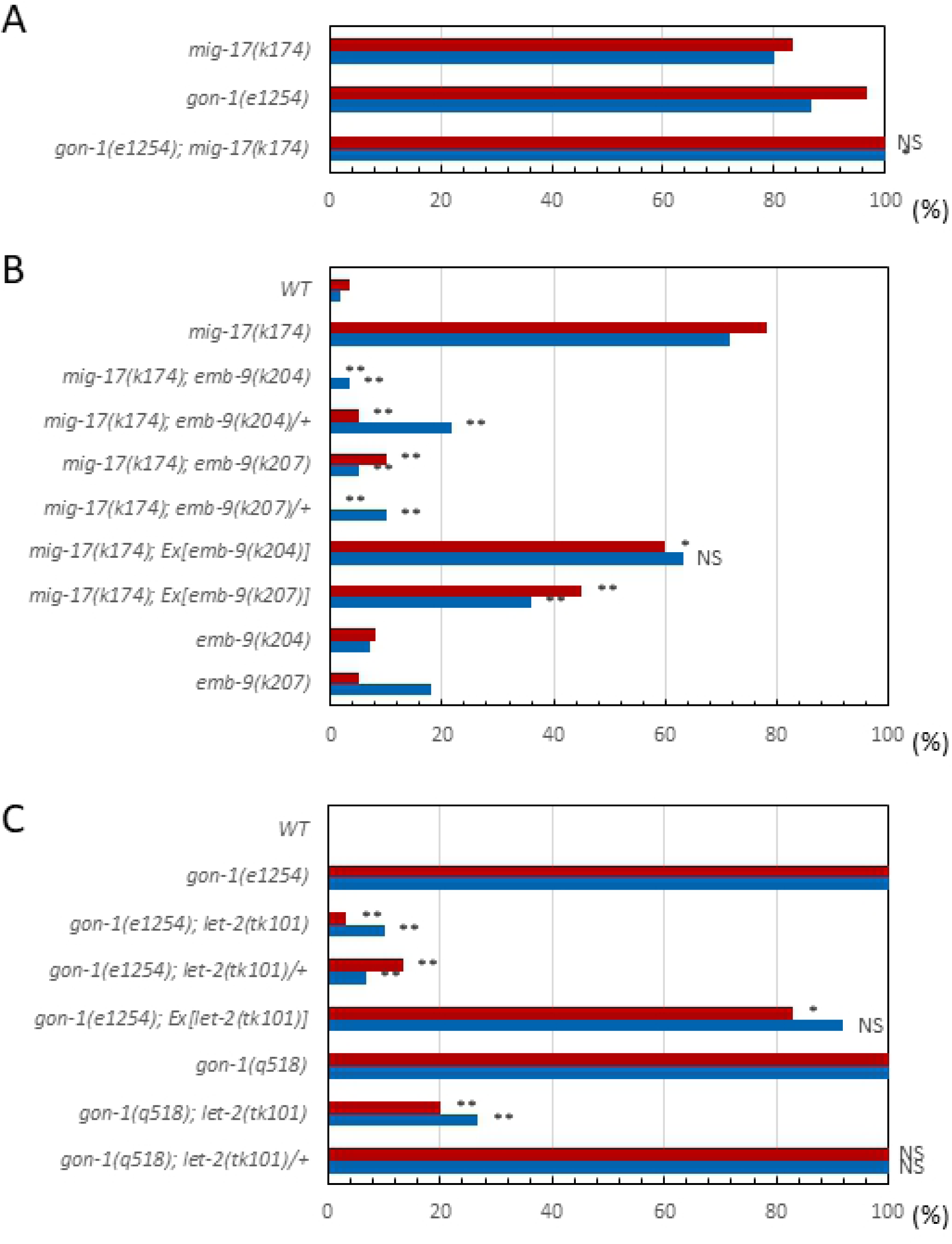
DTC migration phenotypes of *mig-17, gon-1* and the suppressors. **A**. Percentage of DTC migration defects in *mig-17(k174)* and *gon-1(e1254)* single mutants and their double mutants. **B**. Percentage of DTC migration defects in *mig-17(k174)* animals with the *emb-9(k204)* or *emb-9(k207)* mutation or with extrachromosomal arrays carrying these mutant genes. **C**. Percentage of DTC migration defects in *gon-1(e1254)* or *gon-1(q518)* animals with a *let-2(tk101)* mutation or with an extrachromosomal array carrying the *let-2(tk101)* mutant gene.

To identify genes interacting with *mig-17* and *gon-1*, we isolated novel suppressor mutations for DTC migration defects of *mig-17* and *gon-1* mutants using EMS mutagenesis. We isolated 11 suppressor mutants of *mig-17(k174)*, a null allele that has a nonsense mutation in the pro-domain (S2 Fig). Two of these mutants, *k204* and *k207*, were genetically mapped near the center of LG III and acted as dominant suppressors for *mig-17* (Fig 1B). Next-generation sequence analysis of *k204* and *k207* identified mutations in *emb-9*, which encodes the α1 subunit of collagen IV. Both mutations were amino acid substitutions in the C-terminal NC domain. We generated plasmids carrying these mutant alleles of *emb-9* and introduced them into *mig-17* mutants. The extrachromosomal arrays containing these plasmids partially rescued the gonadal phenotype of *mig-17* mutants (Fig 1B), indicating that *emb-9(k204)* and *emb-9(k207)* mutations are causative for *mig-17* suppression.

We used EMS mutagenesis to isolate suppressors of *gon-1(e1254)*, a strong loss-of-function allele with a nonsense mutation in the C-terminal domain, which contains thrombospondin type 1 motifs (S2 Fig). Although *gon-1(e1254)* homozygotes are sterile, the transgenic strain *gon-1(e1254); tkEx370[gon-1 fosmid, rol-6(su1006), sur-5::GFP]*, which carries an extrachromosomal array that consists of multiple copies of the wild-type *gon-1* fosmid (WRM0622dB04), mutant *rol-6(1006)* plasmid and s*ur-5::GFP* plasmid, does proliferate as a homozygote. *rol-6(su1006)* and *sur-5::GFP* are marker plasmids that result in the roller movement phenotype and GFP expression in all somatic nuclei, respectively. We isolated non-roller and GFP^−^ fertile animals from the F_2_ or F_3_ generation of transgenic animals treated with EMS (S3 Fig). One of the two *gon-1* suppressors, *tk101*, acted as a dominant suppressor of *gon-1(e1254)* and was genetically mapped to the right end of LG X. *tk101*was a recessive suppressor for the *gon-1(q518)* null allele (Fig 1C). Next-generation sequence analysis revealed an amino acid substitution in the N-terminal region of the triple helical domain of *let-2*, which encodes the α2 subunit of collagen IV. The extrachromosomal array containing this *let-2* mutant plasmid partially rescued the gonadal phenotype of *gon-1(e1254)* (Fig 1C), indicating that *let-2(tk101)* is the causative mutation for *gon-1* suppression.

### Swapping experiments for *mig-17* and *gon-1* suppressors or enhancers

Thus far, our genetic screening had identified various mutant alleles in *let-2* and *fbl-1* that act as suppressors of *mig-17* [11, 12] and in *fbl-1* that act as a suppressor of *gon-1* [14], among which the suppressor mutant alleles differed between *mig-17* and *gon-1*. We also previously showed that loss-of-function mutations *emb-9(g34* and *g23cg46)* and *let-2(g25* and *b246)* act as suppressors of *mig-17* [12] and that *emb-9(tk75)*, which was originally isolated as a suppressor of *fbl-1(tk45)*, acts as an enhancer of *gon-1* [14]. *emb-9(g34* and *g23cg46)* also enhance *gon-1* [14]. In this study, we identified novel suppressors *let-2(tk101)* for *gon-1* and *emb-9(k204* and *k207)* for *mig-17* (Fig 2). These results imply that MIG-17 and GON-1 ADAMTS proteases functionally interact with the same BM proteins collagen IV α1 and α2 subunits and fibulin-1.

**Fig 2.**
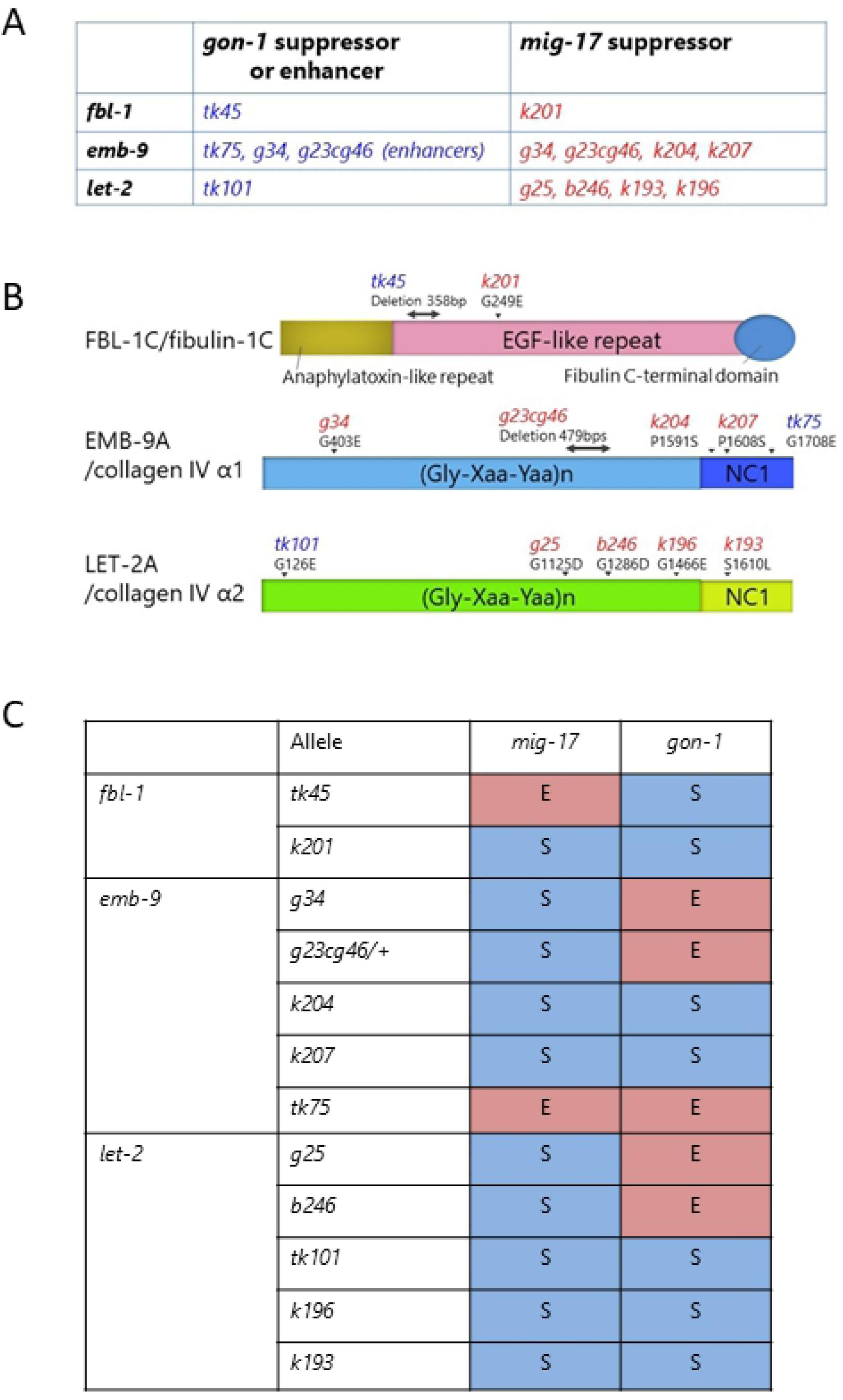
Suppressor and enhancer mutations for *gon-1* and *mig-17*, and summary for swapping experiments of suppressors and enhancers. **A**. *fbl-1, emb-9* and *let-2* alleles that suppress or enhance the gonadal defects of *gon-1(e1254)* or *mig-17(k174)*. **B**. Locations of mutations in FBL-1C, EMB-9A and LET-2A proteins. The mutation sites are shown by arrowheads (amino acid substitutions) or bidirectional arrows (deletions). Both *fbl-1(tk45)* and *emb-9(g23cg46)* deletions are potential null alleles, as they are expected to introduce termination codons shortly after the deleted region [11, 17]. **C**. Summary of effects of *fbl-1, emb-9* and *let-2* alleles on *mig-17(k174)* and *gon-1(e1254)* mutants. S and E represent suppression and enhancement, respectively.

To understand the roles shared by these two proteases, we examined how the suppressors of *mig-17* affect *gon-1* and how the suppressors and enhancers of *gon-1* affect *mig-17* gonadal defects. We examined all the combinations of double mutants. The representative phenotypes exhibited by these double mutants are shown in Fig 3, and their phenotypic penetrance scored at the young adult stage is shown in Fig 4 and Fig 5. In *mig-17(k174)* animals and in double mutants containing *mig-17(k174)*, suppression was assessed by whether the U-shaped gonad was formed as in the wild type. In *gon-1(e1254)* animals and in the double mutants containing *gon-1(e1254)*, suppression was assessed by whether the gonad arms reached the dorsal muscle.

**Fig 3.**
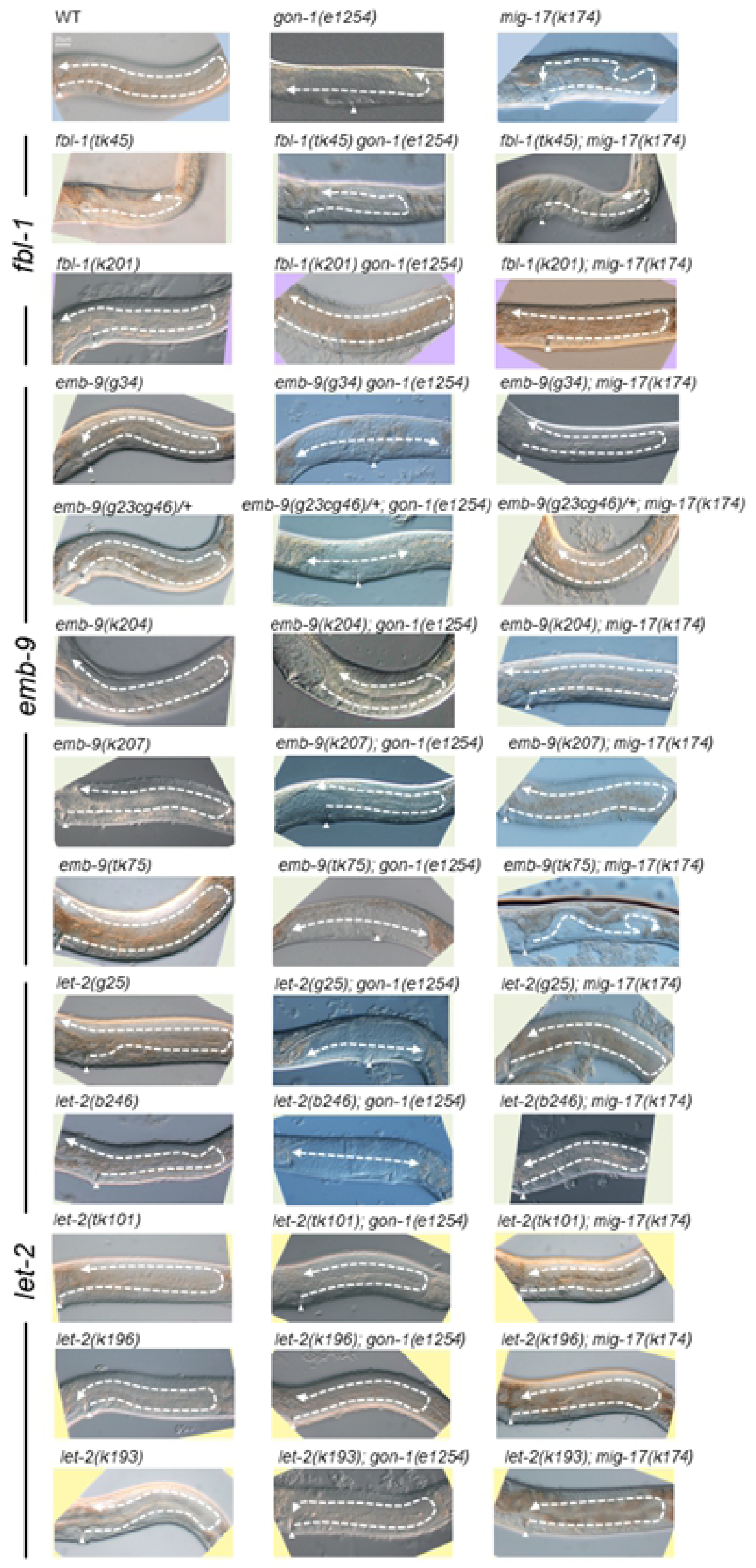
Representative Nomarski images of young adult hermaphrodite gonads of wild-type, *gon-1(e1254), mig-17(k174)* and double-mutant animals analyzed in this study. The gonad morphology is shown by dashed arrows. Anterior is to the left, dorsal up. Arrowheads point to the vulva. Bar: 20 μm.

**Fig 4.**
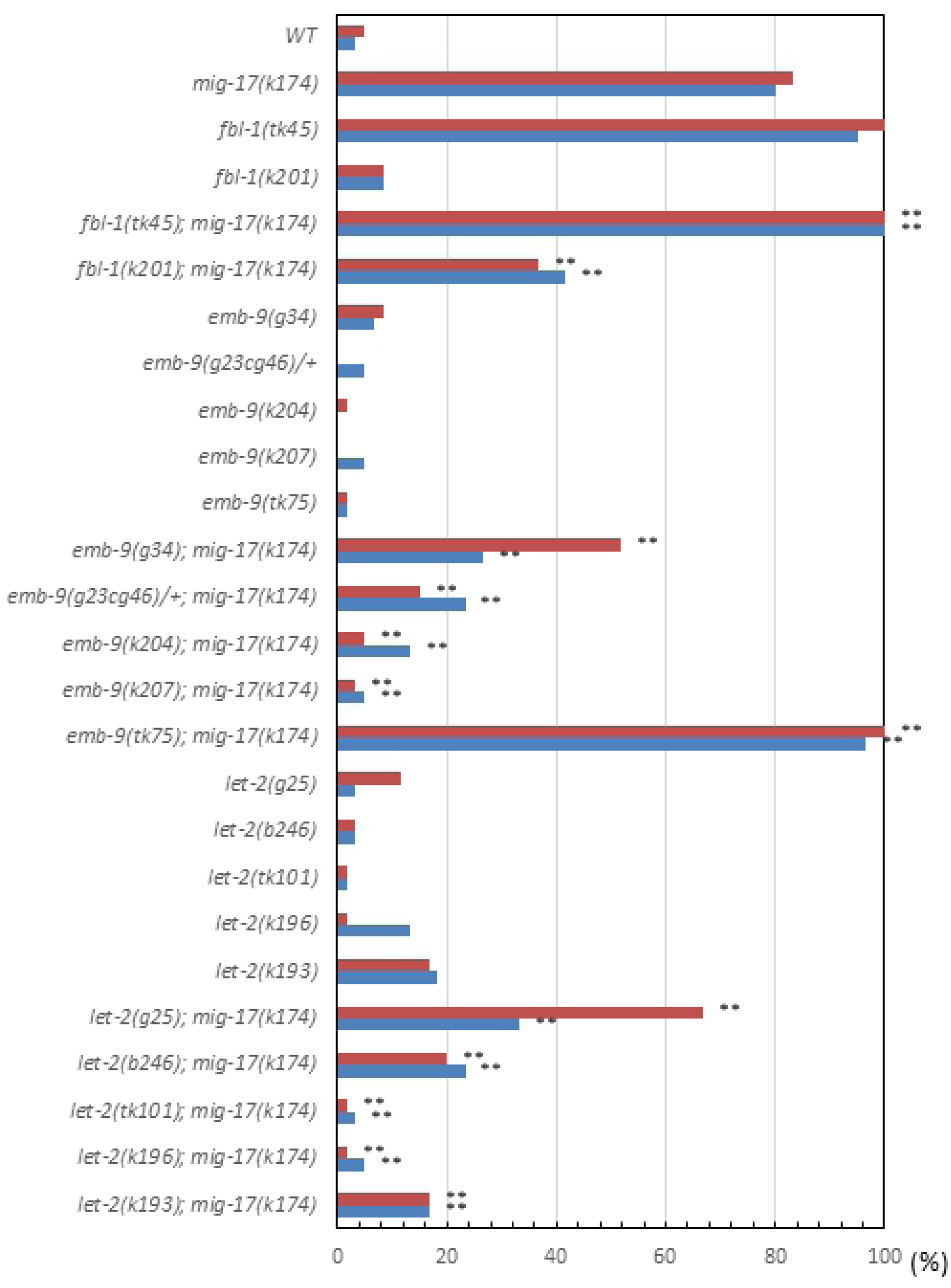
Percentage of DTC migration defects in *mig-17(k174)* mutants in the presence of *fbl-1, emb-9* and *let-2* alleles. *n* = 60 for each experiment. *P*-values from Fisher’s exact test comparing the double mutants with *mig-17(k174)* animals: \*\**P* < 0.01; \**P* < 0.05; NS, not significant.

**Fig 5.**
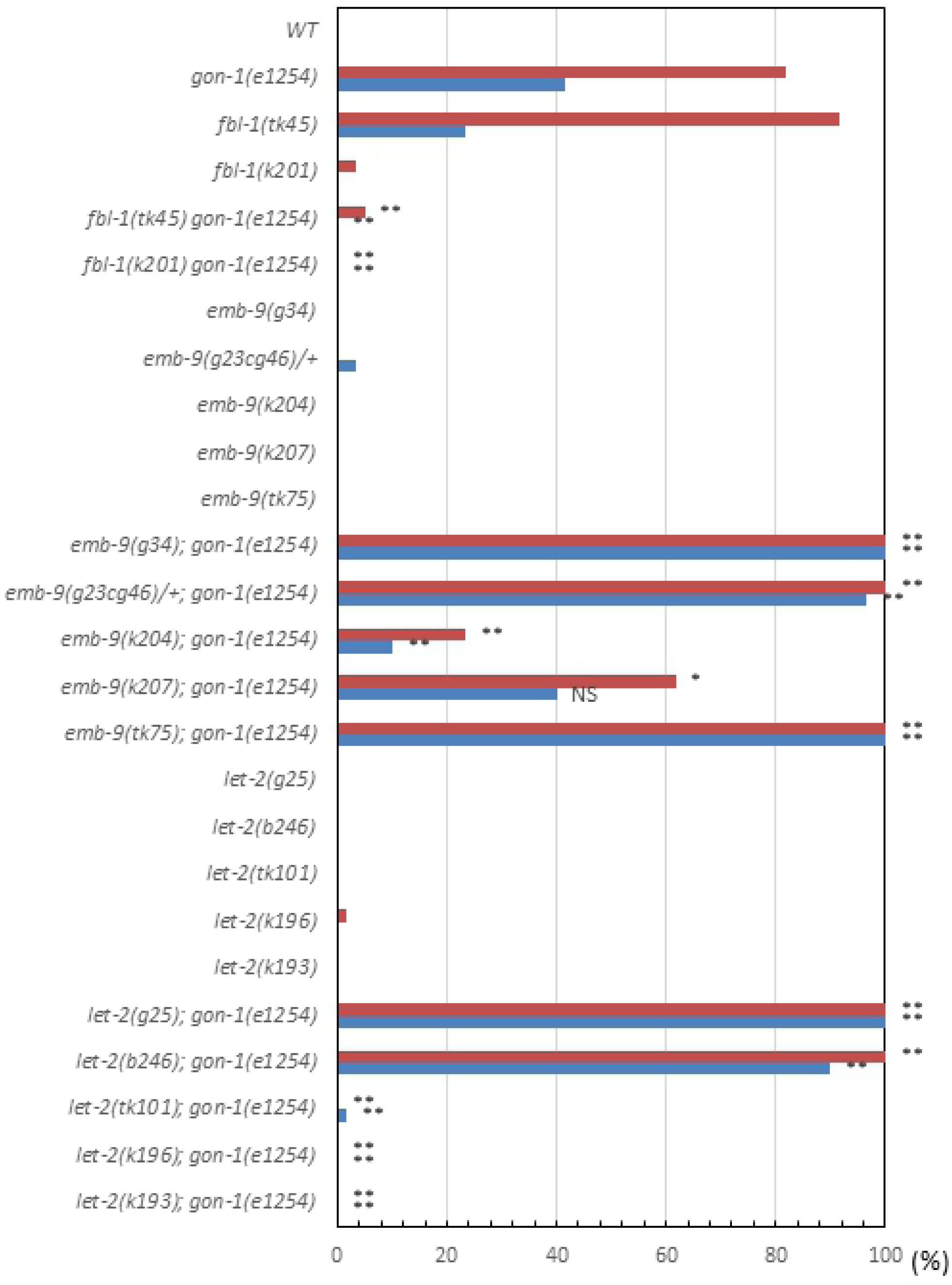
Percentage of gonad arms that failed to reach the dorsal muscle of *gon-1(e1254)* mutants in the presence of *fbl-1, emb-9* and *let-2* alleles. *n* = 60 for each experiment. *P*-values from Fisher’s exact test comparing the double mutants with *gon-1(e1254)* animals: \*\**P* < 0.01; \**P* < 0.05; NS, not significant.

For the *fbl-1* alleles, the *k201* mutation suppressed both *mig-17* and *gon-1*. Although *tk45* suppressed *gon-1*, it rather enhanced *mig-17*. For the *emb-9* alleles, although *g34* and *g23cg46/+* suppressed *mig-17*, they both enhanced *gon-1. k204* and *k207* suppressed *mig-17* strongly, whereas they suppressed *gon-1* somewhat weakly. *tk75* acted as a strong enhancer for both *mig-17* and *gon-1*. For the *let-2* alleles, although *g25* and *b246* suppressed *mig-17*, they both enhanced *gon-1. tk101, k196* and *k193* suppressed both *mig-17* and *gon-1*. These data are summarized in Fig 2C. Because *gon-1* mutants are 100% sterile, we also analyzed fertility in the double mutants (S4 Fig). We found that among *fbl-1, emb-9* and *let-2*, some alleles suppressed or enhanced both *mig-17* and *gon-1*, whereas others affected *mig-17* and *gon-1* differentially. In the latter case, the alleles that suppressed *mig-17* or *gon-1* rather enhanced *gon-1* or *mig-17*, respectively. These results suggested that the former suppressor alleles suppress the common functional defects in *mig-17* and *gon-1*, whereas the latter alleles suppress gene-specific defects in either *mig-17* or *gon-1*.

Three mutations found in EMB-9 (*k204, k207* and *tk75*) and one in LET-2 (*k193*) were localized to the C-terminal NC1 domain of these collagen IV molecules. Using SWISS-MODEL, we deduced the three-dimensional structures of the NC1 domains of EMB-9 and LET-2 based on the crystal structures of bovine collagen IV NC1 domains [24] (Fig 6). We found that three amino acid substitutions in EMB-9, which are separated from one another in the primary sequence, were closely apposed in the three-dimensional structure. In the type IV collagen meshwork, triple-helical collagen molecules connect to one another through NC1-NC1 domain interactions. EMB-9(*k207* and *tk75*) mutations were localized to the NC1-NC1 interface regions, and EMB-9(*k204*) was close to these interface regions, suggesting that these amino acid substitutions may affect physical interactions between two NC1 trimers. However, because these mutants were able to proliferate as homozygotes, their mutant collagen molecules were most likely successfully assembled into the functional network in the BM. It is interesting that the amino acid substitutions EMB-9(*k204* and *k207*), which strongly suppressed *gon-1* and *mig-17*, and EMB-9(*tk75*), which strongly enhanced *gon-1* and *mig-17*, were closely localized in the three-dimensional structure.

**Fig 6.**
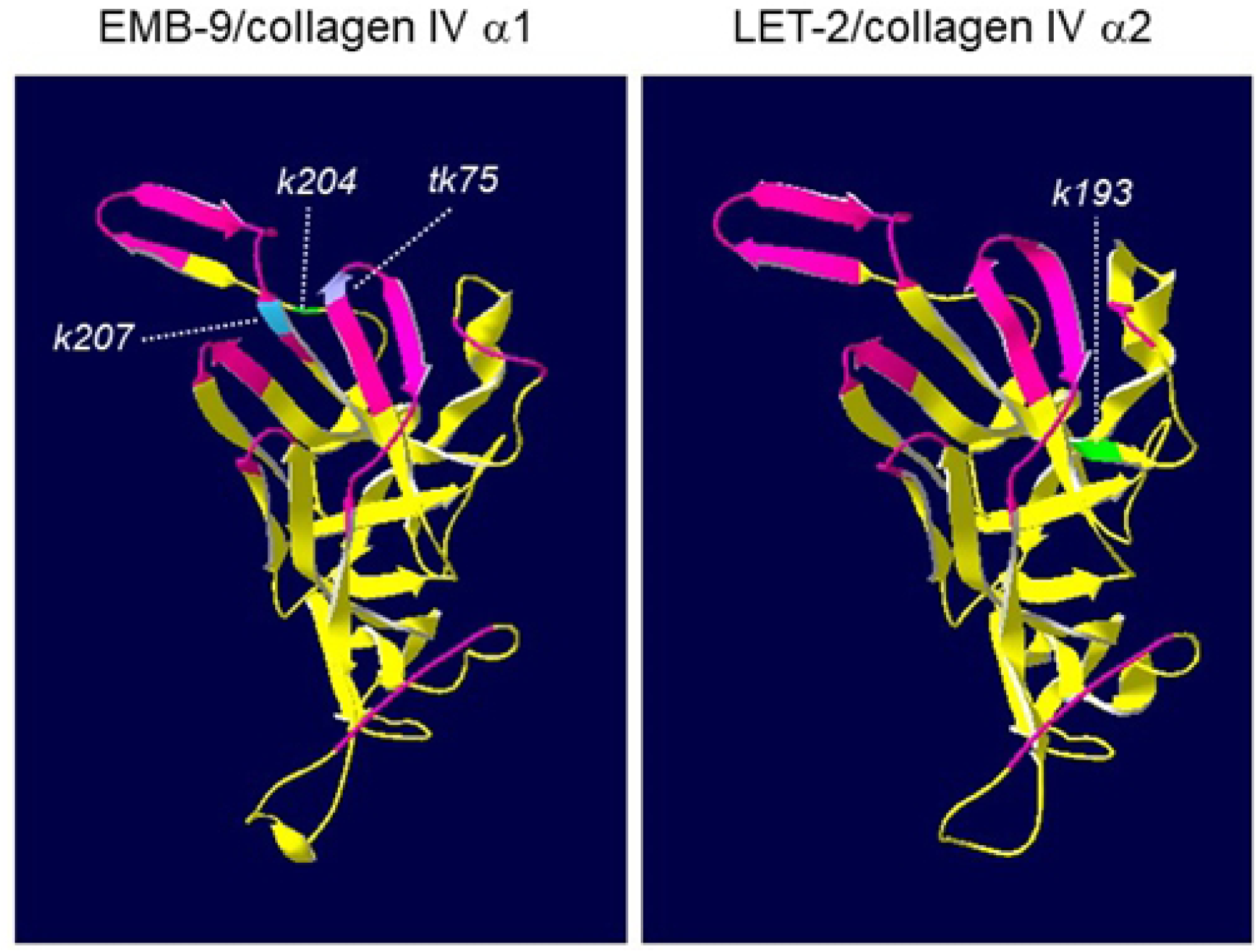
Predicted three-dimensional structures of NC1 domains of *C. elegans* type IV collagen subunits EMB-9 and LET-2. The segments corresponding to the interface region of two NC1 trimers are shown in magenta. The mutated amino acids in the *k204, k207, tk75* and *k193* mutations are highlighted each one in a different color.

### Collagen IV accumulation in the DTC BM

Based on an immunohistochemical analysis, we previously reported that the reduced accumulation of collagen IV in the gonadal BM in animals with *fbl-1(tk45)*, a null mutation, can be compensated by *gon-1(e1254)* [14]. To examine the amount of collagen IV accumulation in the BM quantitatively, we used a functional *emb-9::mCherry* fusion reporter [18]. Third larval stage animals in which their DTCs were at or shortly beyond the first turn were selected, and the fluorescence intensity of the BM surrounding the DTCs was measured (Fig 7A-C). Among the six single mutants examined, we observed that the intensity was slightly higher in *fb-1(tk45)*, significantly lower in *let-2(k196)* and significantly higher in *gon-1(e1254)* as compared with wild type. When combined with *fbl-1(k201* and *tk45)* or *let-2(k196* and *g25)* mutations, the level of EMB-9-mCherry accumulation in *mig-17* mutants was not affected except for the case of *let-2(k196)*, in which a slightly lower accumulation was observed (Fig 7B). Because the three mutations *fbl-1(k201), let-2(k196)* and *let-2(g25)* suppressed *mig-17* but *fbl-1(tk45)* did not, the levels of EMB-9-mCherry accumulation were not correlated with *mig-17* suppression. We then combined *fbl-1(k201* and *tk45)* or *let-2(k196* and *g25)* mutations with *gon-1*. We observed that the levels of EMB-9-mCherry accumulation in *fbl-1(k201), fbl-1(tk45)* and *let-2(k196)*, all of which suppress *gon-1*, were significantly lower than that of the *gon-1* single mutants, whereas the level was not affected in *let-2(g25)*, which enhances *gon-1* (Fig 7C). Thus, it is possible that the reduction in EMB-9-mCherry accumulation is indicative of *gon-1* suppression.

**Fig 7.**
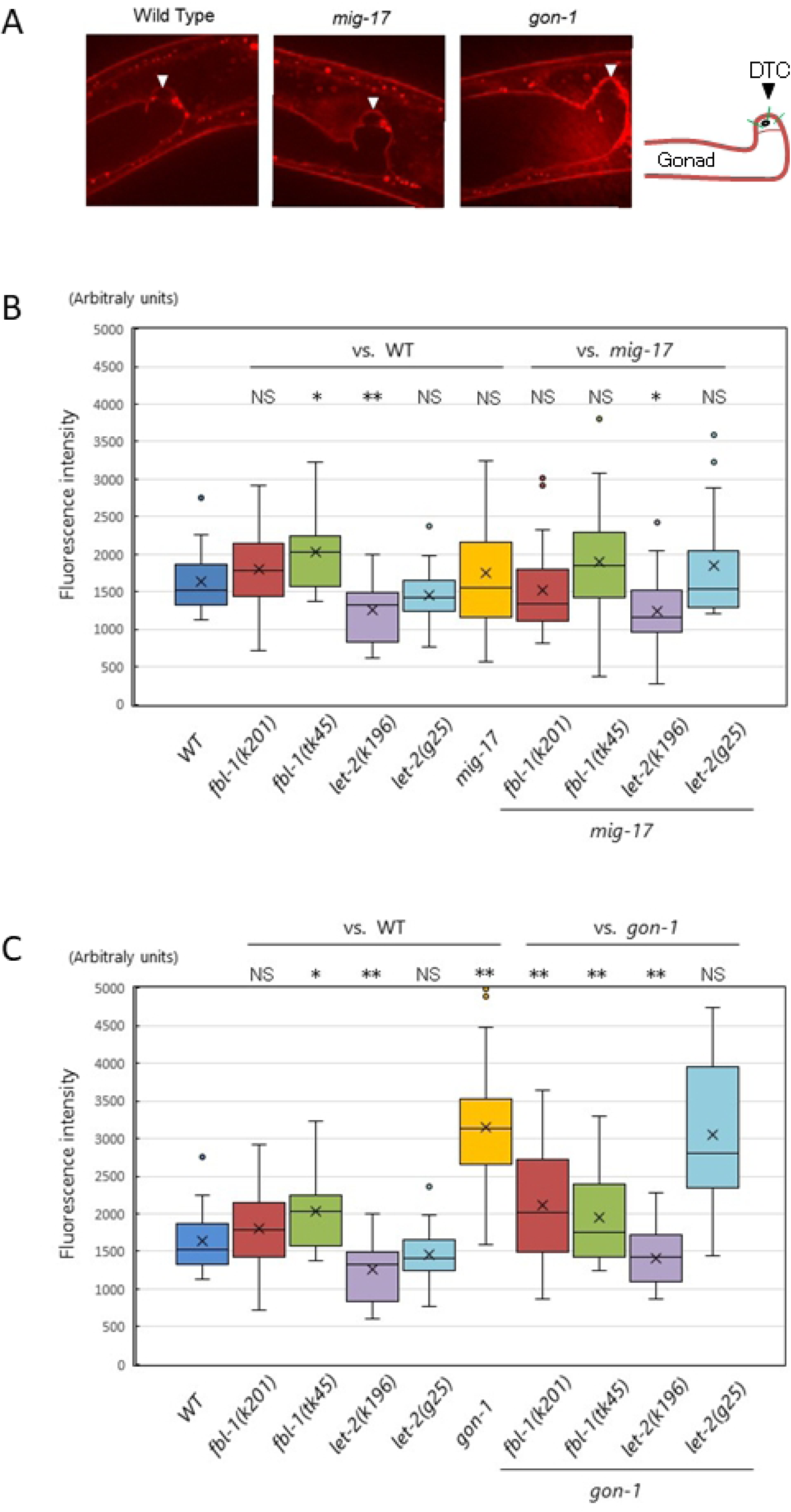
Accumulation of emb-9-mCherry in the BM. **A**. Representative images of optical sections of the gonadal tip shortly after the first turn of the DTCs in wild-type, *mig-17(k174)* and *gon-1(e1254)* animals expressing EMB-9-mCherry. Arrowheads point to DTCs. The right panel illustrates the gonadal BM (brown). The fluorescence intensity of the DTC BM was quantified by averaging the peak values along three lines (green) that cross the DTC BM (see Materials and methods). **B**., **C**. Box-and-whisker plot of the fluorescence intensity of EMB-9-mCherry in the DTC BM in animals with wild-type and mutant *fbl-1* and *let-2* alleles and with those mutant alleles in combination with *mig-17(k174)* (B) or *gon-1(e1254)* (C); *n* = 20. Boxplots indicate the median and the interquartile range. Whiskers extend to the minimum and maximum values within 1.5 times the interquartile range. Points indicate outliers. *P*-values from Student’s t-test are indicated: \*\**P* < 0.01; \**P* < 0.05; NS, not significant.

## Discussion

In the present study, we isolated novel suppressor mutations of gonadal defects related to *mig-17* and *gon-1* mutants. We identified alleles of *emb-9* as *mig-17* suppressors and an allele of *let-2* as a *gon-1* suppressor for the first time. We found that some of the *emb-9 (collagen IV a1), let-2 (collagen IV a2)* and *fbl-1* (*fibulin-1*) mutations suppressed both *gon-1* and *mig-17*, whereas others suppressed only *gon-1* or *mig-17*. These results suggest that *gon-1* and *mig-17* have their specific functions as well as functions in common that relate to gonad formation. Probably, the loss of the gene-specific functions is the cause of the very different phenotypes of the *gon-1* and *mig-17* mutants.

The *fbl-1(tk45)* null mutation suppressed *gon-1* but enhanced *mig-17*. The *gon-1* suppression is likely to be due to the reduction of collagen levels in the late larval stages that results from loss of *fbl-1* activity [14], although the collagen levels were not reduced in the mid-L3 stage when the DTCs make their first turn (Fig 7C). FBL-1C acts downstream of MIG-17 to recruit NID-1/nidogen-1 to regulate directed DTC migration [12]. Thus, *mig-17* is enhanced probably because of the reduction of NID-1 in the DTC BM. In *mig-17* mutants, the DTCs do not migrate along their normal U-shaped route because they often detach from the body wall (their normal migratory substratum) and mis-attach to the intestine [8]. Therefore, NID-1 is likely to be required for appropriate adhesiveness between the DTC and body wall BMs. The *fbl-1(k201)* gain-of-function mutation suppressed both *mig-17* and *gon-1. fbl-1(k201)* suppressed the collagen accumulation in *gon-1*, as did *fbl-1(tk45)*, but *fbl-1(k201)* is fully fertile on its own, unlike *fbl-1(tk45)* (S3 Fig). Therefore, it is possible that the *fbl-1(k201)* mutation may partially compromise the ability of FBL-1C to maintain collagen IV without affecting its ability to recruit NID-1.

The gain-of-function mutations of collagen IV *emb-9(k204, k207)* and *let-2(tk101, k196* and *k193)* were potent suppressors of *mig-17* and *gon-1*. We found that the BM collagen levels in *let-2(k196)* were significantly decreased and that *let-2(k196)* suppressed the dramatic increase in collagen accumulation in *gon-1* animals. Although we did not examine the other gain-of-function collagen mutations, it is possible that the levels of BM collagen are similarly affected. In contrast, the gain-of-function mutation *emb-9(tk75)* enhanced both *mig-17* and *gon-1*. We previously showed that the levels of BM collagen in animals expressing the mutant EMB-9(*tk75*) α1 subunit can be maintained in the absence of FBL-1, which is otherwise required for the maintenance of BM collagen levels [14]. Therefore, it is likely that too much accumulation of collagen in the BM could be causative for both the *mig-17* and *gon-1* mutant phenotypes. Although we could not detect over-accumulation of collagen in the *mig-17* mutant, this could be because of the insufficient sensitivity of our assay condition.

This model is not, however, consistent with the fact that the loss-of-function mutations of collagen IV, *emb-9(g34* and *g23cg46/+)* and *let-2(g25* and *b246)*, which conceivably reduce the levels of BM collagen, all acted as enhancers of *gon-1* even though they acted as suppressors of *mig-17*. The loss-of-function *emb-9(xd51)* mutation also enhances *gon-1* in the presynaptic bouton overgrowth phenotype [25]. In contrast, *emb-9(g34)* and *emb-9(b189)* can suppress the synaptic defect of *gon-1* when *emb-9; gon-1* double mutants are shifted up from 16 to 25°C after completion of embryogenesis [26]. In this case, it is possible that collagen IV containing the temperature-sensitive mutant proteins EMB-9(*g34*) and EMB-9(*b189*) form intracellular aggregates and thus are not secreted [16], and therefore the BM accumulation of collagen could be considerably decreased. Our analysis using the EMB-9-mCherry reporter revealed that the collagen levels of *let-2(g25); gon-1* or *let-2(g25); mig-17* double mutants were not lowered relative to the respective *gon-1* or *mig-17* single mutant (Fig 7B, C). It might be possible that a subtle reduction in collagen accumulation is sufficient for amelioration of BM physiology in *mig-17* mutants but instead worsens that in *gon-1* mutants. The slight collagen reduction may promote NID-1 accumulation in the BM and suppress the *mig-17* gonadal defect.

Why is *gon-1* enhanced by collagen IV loss-of-function mutations? This seemingly contradictory phenomenon may be related to the dual function of GON-1. In addition to its predicted extracellular protease activity, GON-1 functions in the endoplasmic reticulum to transport secreted or membrane proteins from the endoplasmic reticulum to the Golgi. This transport activity is dependent on the C-terminal GON domain but is independent of protease activity [27]. For example, cell surface expression of the integrin receptors that are required for cell migration may be reduced and, therefore, the DTC migration activity in *gon-1* mutants may be weakened. Because the remodeling of the BM is coupled with epithelial cell migration [28, 29], reduced migration of DTCs could also lead to downregulation of BM remodeling, resulting in thick accumulation of collagen, which can further block DTC migration. Remodeling of the BM is likely to be mediated by the GON-1 protease activity.

We previously showed that a reduction in collagen IV lowers PAT-3/β-integrin expression in DTCs [14]. At the early second larval stage, the fluorescence levels of EMB-9-mCherry in *gon-1* mutants were closer to those in the wild type (K.N., unpublished). Thus, it might be possible that when combined with loss-of-function collagen IV mutants, the reduced collagen levels may further impair integrin expression in DTCs of *gon-1* animals in the second larval or younger stages. This might be the reason for the enhancement of the *gon-1* phenotype in the presence of collagen IV loss-of-function mutations.

It is unclear why the gain-of-function collagen IV mutation *let-2(k196)* did not enhance *gon-1*, as did *let-2(g25)*, even though *let-2(k196)* also reduced collagen accumulation. We speculate that GON-1 is the major enzyme responsible for turnover of BM collagen IV and that the mutant LET(*k196*) protein may confer the property by which the collagen IV meshwork is turned over quickly as compared with the wild-type meshwork even with the weakened activity of GON-1(*e1254*) mutant enzyme. If this is the case, the collagen meshwork containing EMB-9(*tk75*) may have gained a slower turnover rate.

Our observations suggest that both MIG-17 and GON-1 function to reduce collagen IV in the BM. Although this is consistent with the idea that collagen IV is the direct substrate of these enzymes, we still do not have evidence of this interaction. Future biochemical analysis is thus needed to determine the substrates. In addition, molecular structural analysis of interactions among these ADAMTS proteases and the BM molecules should shed light on the detailed mechanism of BM remodeling during organogenesis.

## Acknowledgments

We thank Noriko Nakagawa and Nami Okahashi for technical assistance. Some nematode strains used in this work were provided by the Caenorhabditis Genetics Center, which is funded by the National Institutes of Health National Center for Research Resources. This work was supported by a Grant-in-Aid for Scientific Research on Innovative Areas by Ministry of Education, Culture, Sports, Science and Technology to KN (22111005) and by the Naito Grant for the advancement of natural science to KN.

## Supporting information

**S1 Fig. Schematic presentation of gonad formation in the *C. elegans* hermaphrodite**. The hermaphrodite U-shaped gonad arms are generated by migration of two DTCs. DTCs are generated at the anterior (A) and posterior (P) ends of the gonad primordium at the first larval stage (L1) and migrate along the anteroposterior axis on the ventral (V) body wall muscle (L2–L3). The DTCs turn dorsally (D) and migrate along the lateral hypodermis (L3). Finally, the DTCs undergo a second turn and migrate along the dorsal body wall muscle to form the symmetrical U-shaped arms (L4).

**S2 Fig. Structure and mutation sites of GON-1 and MIG-17 proteins**. Domain organization is shown by colored boxes. Mutation sites for *gon-1(q518* and *e1254)* and *mig-17(k174)* are indicated.

**S3 Fig. Isolation of *gon-1* suppressors**. Gonad arms (arrows) of young adult hermaphrodites are shown. **A**., **B**. Nomarski images of wild type (A) and *gon-1(e1254)* (B). **C**., **D**. Nomarski (C) and fluorescence (D) images of a *gon-1(e1254); tkEx370[gon-1 fosmid, rol-6(su1006), sur-5::GFP]* young adult hermaphrodite. **E**., **F**. Nomarski (E) and fluorescence (F) images of a *let-2(k101); gon-1(e1254)* young adult hermaphrodite, which lost the *tkEx370[gon-1 fosmid, rol-6(su1006), sur-5::GFP]* extrachromosomal array.

**S4 Fig. Percentage of fertile animals**. For each strain, 20 first larval stage hermaphrodites were grown at 20°C, and the number of animals that produced offspring were counted. Green and red bars represent % fertile and % sterile animals, respectively.

